# Behavioral evidence for feedback gain control by the inhibitory microcircuit of the substantia nigra

**DOI:** 10.1101/090209

**Authors:** Jennifer Brown, Kathleen A. Martin, Joshua T. Dudman

## Abstract

We recently demonstrated that the collaterals of substantia nigra (SN) projection neurons can implement divisive feedback inhibition, or gain control (Brown et al., 2014). While in vivo recordings were consistent with divisive feedback inhibition, a causal test was lacking. A gain control model of the nigral microcircuit implies behavioral effects of disrupting intranigral inhibition that are distinct from previous functional models. To test the model predictions experimentally, we develop a chemogenetic approach that can selectively suppress synaptic release within the substantial nigra without affecting the propagation of activity to extranigral targets. We observe behavioral consequences of suppressing intranigral inhibition that are uniquely consistent with a gain control model. Our data further suggest that if endogenous metabotropic signaling can modulate intranigral synapses, this would provide a circuit mechanism for an exploitation/exploration trade-off in which the timing and variability of goal-directed movements are controlled independently of changes in action.

## Introduction

The canonical basal ganglia circuit is composed of nuclei that project, in largely feed-forward fashion, from input nuclei, striatum and subthalamic nucleus (STN), to output nuclei, substantia nigra pars reticulata (SNr) and internal globus pallidus (GPi) (Dudman and Gerfen, 2015). From input to output, the number of projection neurons decreases dramatically from a few million in the striatum to roughly thirty thousand in the SNr (Oorschot, 1996). The reduction in number of neurons dedicated to the representation of corticothalamic output implies either a loss of information (Dudman and Krakauer, 2016) or a mechanism for a compact representation (Bar-Gad et al., 2003). In many circuits, local interneurons have been shown to perform important computational roles such as lateral inhibition or divisive gain control on projection neuron outputs (Isaacson and Scanziani, 2011). While processing of activity by local microcircuits in the basal ganglia is attractive, a low percentage of total striatal neurons are local interneurons and none have been found, to date, in the SNr (Deniau et al., 2007). However, projection neurons in the SNr, which are tonically active and inhibitory, elaborate axon collaterals within the SNr (intranigral) and we have recently argued (Brown et al., 2014) may perform computational roles akin of interneurons in cortical microcircuits (Silver, 2010).

The idea that the basal ganglia performs action selection by producing sparse, selective representations of actions (Mink, 1996) (in a compact form (Bar-Gad et al., 2003)) leads to the proposal that intranigral collateral inhibition could implement lateral inhibition (Deniau et al., 2007). However, several lines of evidence suggest that the basal ganglia are also critical for the control of movement execution (Dudman and Krakauer, 2016; Turner and Desmurget, 2010), specifically in mice (Panigrahi et al., 2015; Yttri and Dudman, 2016). This suggests that continuous and stable dynamics of activity during the multiple phases of movement execution are also critical (Dudman and Krakauer, 2016). This latter possibility implies a need for some form of gain control (a hallmark of engineered control systems) at the basal ganglia output instead of, or perhaps in addition to, lateral inhibition. Our prior work found little evidence of a functional organization in the SN consistent with lateral inhibition (Brown et al., 2014). However, we argued that intranigral collaterals, when combined with the intrinsic biophysical properties of projection neurons, are sufficient to implement divisive feedback inhibition - a form of gain control (Brown et al., 2014). At the time, we provided evidence that intranigral inhibition was sufficient to produce robust feedback inhibition *in vivo*. Moreover, recordings of population activity in the SNr in behaving mice exhibited noise correlations consistent with gain control (Brown et al., 2014). However, a causal test - perturbing intranigral inhibition and observing its consequences on the control of purposive behaviors - was lacking.

The critical experimental need was to interfere with intranigral synaptic transmission without altering the spiking of nigral projection neurons (and thereby disrupting propagating activity to extranigral targets). Here we show that pharmacogenetic activation of Gi-coupled signaling (using the hM4Di designer receptor activated by designer drug [DREADD] receptor (Armbruster et al., 2007)) profoundly suppresses synaptic release in SNr projection neurons but has no measurable effect on somatic excitability. Recent work demonstrated the potency of hM4Di receptors in suppressing synaptic transmission largely independent of the suppression of a cell’s ability to fire action potentials (Stachniak et al., 2014). In SNr projection neurons, hM4Di-mediated signaling appears specific to synaptic release - alleviating a requirement to bias receptor expression to the axon (Stachniak et al., 2014). Combining this approach with either targeted local infusion of the hM4Di agonist clozapine-N-oxide (CNO) into the SNr or systemic administration, we could critically evaluate whether disrupted intranigral inhibition is consistent with altered gain control in behaving mice. Our data provide the first causal evidence that disruption of intranigral inhibition is consistent with a feedback gain control model and are inconsistent with other prominent models in which intranigral inhibition is proposed to implement lateral inhibition (Deniau et al., 2007; Mink, 1996) or tonic suppression of output (Chevalier and Deniau, 1990).

## Results and Discussion

We previously argued that to produce divisive feedback inhibition, a specific combination of diverse tonic firing rates, spatially-diffuse collateral circuit, and nonlinear subthreshold response to asynchronous inhibition was sufficient (Brown et al., 2014). Asynchronous, weak inhibition amongst reciprocally coupled, tonically active neurons requires a low rate of pairwise connectivity and relatively weak unitary connections. We provided evidence that these properties are present in the intranigral microcircuit (Brown et al., 2014). However, as we noted in our discussion, it was difficult to envisage how to test such a model in a behaving mouse. A major confound to experimentally overcome in a microcircuit like the SNr is that somatic spiking and synaptic release are spatially intermingled. Thus, manipulation of excitability locally at the site of synaptic release in the axon collateral is difficult to disentangle from a manipulation of nearby somatic excitability. Subsequently, it became clear that in some cells, exogenously controlled Gi-coupled signaling can be sufficient to dramatically reduce the release of vesicles from synaptic terminals through a distinct mechanism from the modulation of somatic excitability or spike propagation (Stachniak et al., 2014). Thus, here we took advantage of this unique experimental approach to ask whether suppressing feedback inhibition locally within the SNr is consistent with a disruption of feedback gain control of the basal ganglia output.

While the DREADD receptor hM4di makes this experiment (in principle) feasible, a challenging issue remains: the precise role of the basal ganglia in behavior is still under intense debate, leaving the effect of feedback gain control on behavior as an open question. To compare amongst functional models we note that previous models must at least make internally consistent predictions. This allows us to produce a comparison set of behavioral predictions for functional models in which intranigral inhibition mediates either an unstructured, tonic inhibitory tone or lateral inhibition ((Table 1). Each sign is inferred relative to the sign of the effect of systemic suppression of nigral output (assumed to be the same for all models).

**Table 1.** *Comparison between predictions of three functional models of the nigral microcircuit and experimental observations.*

| Movement Parameter | TONIC<br>(Chevalier & Deniau, 1990) |  | LATERAL<br>(Mink, 1996) |  | GAIN<br>(Brown, et al, 2014) |  | Data |  |
| --- | --- | --- | --- | --- | --- | --- | --- | --- |
|  | Systemic | Local | Systemic | Local | Systemic | Local | Systemic | Local |
| Mean vigor | ↑↑ | ↓↓ | ↑↑ | ↓ | ↑↑ | • | ↑↑ | • |
| Peak velocity | ↑ | ↓ | ↑ | ↓ | ↑ | ↑ | ↑ | ↑ |
| Movement initiation | ↓/• | ↑ | ↓/• | ↑ | ↓/• | ↓ | ↓/• | ↓ |

There are three broad classes of qualitative models attributing specific functions to the collateral connectivity within the SNr. Perhaps the simplest model is one in which intranigral connectivity is simply a source of tonic inhibition that determines how readily disinhibition (and thereby release of a movement) will occur (Chevalier and Deniau, 1990). Second, as focus shifted to the importance of the basal ganglia in the selection of movements, authors began to argue that collateral synapses in the striatum and SNr might implement a form of lateral inhibition that mediated competition between motor programs (Deniau et al., 2007; Mink, 1996). Finally, we consider the model that we recently proposed - that intranigral collaterals mediate a feedback gain control on the SNr output (Brown et al., 2014). In the latter case, gain control, some predictions can appear counterintuitive and thus we also developed a computational model to validate the qualitative predictions.

We first consider ‘movement vigor’ in this case, indexed by the average amplitude of movement (Dudman and Krakauer, 2016). In all models, the systemic suppression of tonic, inhibitory nigral output, which normally constrains the vigor of movement execution, should produce an increase in the vigor of movements ((Table 1; row 1). However, the three proposed functional models of intranigral inhibition make distinct predictions. For the tonic inhibition (‘tonic’) model, if intranigral inhibition tends to tonically reduce inhibitory output to extranigral targets, then local suppression of intranigral inhibition should produce a change in movement vigor that is opposite in sign to systemic suppression. If intranigral inhibition mediates lateral inhibition (‘lateral’), then we should again expect to observe a change in the sign of the behavioral effect between systemic and local suppression of nigral inhibition due to increased competition amongst motor programs (the logic of this prediction was laid out in (Mink, 1996)1). In neither case do the ‘tonic’ nor ‘lateral’ models make specific predictions about the variance of movement velocity (Chevalier and Deniau, 1990; Mink, 1996). However, a correlated change in variance and mean is a general principle of motor control due to execution noise downstream of selection (Bays and Wolpert, 2006). In contrast to the previous models, a change in the gain (‘gain’, (Table 1) of the nigral output (i.e. slope, but not offset (Brown et al., 2014)) is expected to produce little to no change in average amplitude of movements since amplitude results from the integration of positive and negative modulation of instantaneous velocity/acceleration commands (Dudman and Krakauer, 2016; Panigrahi et al., 2015). However, the gain model does predict an increase in the peak velocity since, by construction, such a measurement depends only upon the change in maxima of the input-output function that result from a change in gain (Brown et al., 2014; Silver, 2010).

Lastly, we consider the timing of movement initiation. According to all 3 models systemic suppression of nigral output would be expected to, if anything, decrease the latency to movement initiation by reducing the tonic inhibition of movement (Hikosaka and Wurtz, 1985). However, it is important to note that previous work with ‘systemic’ perturbations found no effect on movement initiation for cued reaching movements (Desmurget and Turner, 2010; Horak and Anderson, 1984). This conflicting experimental data is indicated by a uncertain prediction 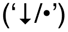 in (Table 1. Again however, the tonic and lateral inhibition models predict a sign change when local intranigral inhibition is blocked compared with systemic suppression. In the case of the lateral inhibition model the reduced contrast between intended and non-intended motor representations predicts a slowing of initiation^1^. In contrast to both tonic and lateral models, the gain control model predicts a decrease in latency to initiate movement following intranigral suppression since enhanced gain would, in fact, *enhance* the magnitude of transient signals that are presumably related to movement initiation.

The predictions of our gain control model were, at least for the authors, counterintuitive at first. Thus, we sought to confirm the predictions of a gain control model by implementing a computational model of the basal ganglia circuit. The SNr circuit model consists of integrate-and-fire units with a nonlinear bias current (Fig. 1A-B; ‘bIF’) that we previously showed to be critical for rendering SNr projection neurons relatively insensitive to perithreshold, asynchronous inhibition (Brown et al., 2014). The properties of the bIF units were drawn from a broad distribution (Fig. 1B) to approximate the distribution of firing rates in the SNr (Brown et al., 2014; Pan et al., 2013). As predicted in (Brown et al., 2014), a network of such units when coupled with relatively sparse intrinsic connectivity (see Methods), produces a robust feedback gain control on spiking output (Fig. 1C).

**Figure 1.**
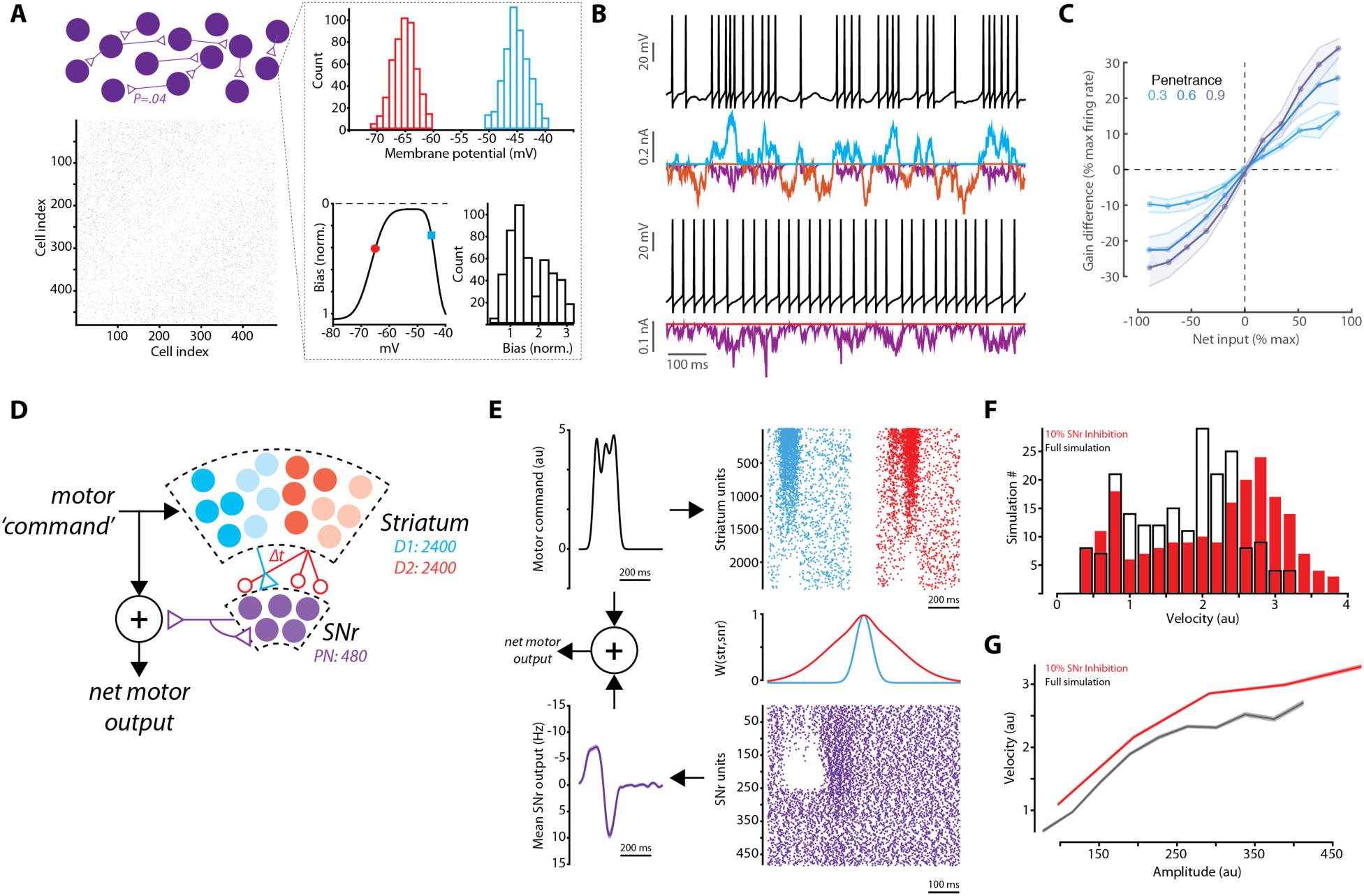
**Intranigral inhibition regulates the gain of the basal ganglia output** **A.** Description of the computational model of the SNr. experiment. Individual integrate-and-fire units were implemented with a nonlinear bias current (bIF). Spike threshold (cyan) and spike reset potential (red) were randomly drawn for each unit (distributions in the simulation used for the figure are shown in histograms). The magnitude of the bias current was also drawn from a random distribution with 2 modes to produce a broad distribution of intrinsic firing rates. **B.** Two example SNr bIF units with similar bias current amplitudes are shown. One unit is driven by coherent external input (upper, red and cyan currents) and the other receives no extrinsic input (lower). Currents derived from the local intranigral circuit are shown in purple. Intrinsic feedback input acts to cancel excitatory extrinsic input (cyan) and a disinhibition can amplify inhibitory (red) extrinsic inputs. **C.** The gain of the network is represented as the percent change in net spike count across the simulated SNr network. The gain effect is proportional to the extent to which intranigral inhibition is removed (‘Penetrance’). **D.** Schematic of the reduced basal ganglia circuit simulation used to formulate predictions about the behavioral consequences of interfering with intranigral inhibition. The SNr network was implemented as in **A.** Multiphasic motor commands were generated and used to produce a distributed pattern of activity in the direct (cyan) and indirect (red) pathways from striatum to SNr. A relatively broad, sign reveresed, and delayed input to SNr from indirect pathway was assumed. **E.** Example simulation data of the reduced basal ganglia circuit simulation from multiphase motor command (top left) through to SNr output (purple, bottom left) is shown. **F-G.** Simulations were run for 200 independently generated motor commands and the predicted motor output is shown (F. peak velocity and G. peak velocity vs. amplitude) for simulations with intranigral inhibition intact (black) and simulations in which 90% (0.9 penetrance) of connections were reduced to 2% of their weights (red).

Recent studies have indicated that activity in the basal ganglia is necessary and sufficient to control the vigor of a purposive movement of the forelimb in mice (Panigrahi et al., 2015; Yttri and Dudman, 2016). Inactivation of specific pathway components can alter movement kinematics (Desmurget and Turner, 2010; Panigrahi et al., 2015), whereas perturbation of all activity in an upstream nucleus (striatum) increases the variance of movement (Rueda-Orozco and Robbe, 2015). These data can be explained by a functional circuit model in which the basal ganglia pathways act like a parallel circuit that can re-converge with motor commands in premotor targets to alter movement vigor (Dudman and Krakauer, 2016; Yttri and Dudman, 2016). In a previous, simplified model of this architecture (Yttri and Dudman, 2016) we did not include the SNr. Thus, here we enrich that prior model with the circuit model of the SNr described in Fig. 1A.

Briefly, we implemented a model of basal ganglia as a parallel circuit that receives copies of motor commands as input to striatum. We simplified the internal connectivity of basal ganglia (Dudman and Gerfen, 2015) such that the indirect pathway produced a spatially broad and temporally delayed excitatory input to the SNr network similar to previous models (Humphries et al., 2006). The SNr output downstream from direct pathway input was integrated at ‘premotor’ structures with motor commands directly from cortex to produce a predicted, net motor output (Yttri and Dudman, 2016). We note that such a functional architecture is consistent with an extensive literature on both normal and pathological basal ganglia function (Dudman and Krakauer, 2016). To test the model across a behaviorally-relevant range of simulated movements, we assumed a variable, multiphase motor command signal that varied in its ratio of direct and indirect striatal units recruited (Fig. 1D). These motor commands were similar to complex movement kinematics observed in a forelimb reaching task we recently described (Panigrahi et al., 2015; Yttri and Dudman, 2016). A substantial fraction (∼40%) of both direct and indirect pathway units in the striatum were positively correlated with the motor command consistent with recent data (Cui et al., 2013; Panigrahi et al., 2015; Yttri and Dudman, 2016).

The model described above makes clear predictions about how the kinematics of a motor command would be altered by suppressing intranigral inhibition. Specifically, the model predicts an increase in the variance (positive skew) of the peak movement velocity (Fig. 1F). Interestingly, the integrated effect of a change in gain (*e.g.* amplitude) is predicted to be relatively unchanged. The lack of amplitude change, despite increases in velocity variance, is a consequence of a divisive gain effect (in contrast to an additive gain) (Silver, 2010). In other words, an increase in gain both enhances the suppression during net inhibition and increases activity during net excitation (Brown et al., 2014). As a result the max speed as a function of total amplitude increases (Fig. 1G). Finally, movement initiation, although not explicitly modeled here, is predicted to change by an analogous argument. Disinhibition below a threshold is thought to be related to movement initiation (Chevalier and Deniau, 1990) and thus, by enhancing the suppression of spiking an increase in gain can lead to earlier movement initiation given the same activity during preparation (summarized in (Table 1).

The functional model of the basal ganglia circuit described above is both consistent with a number of previous observations from our own work in mice (Panigrahi et al., 2015; Yttri and Dudman, 2016) as well as that in other mammals (Anderson and Horak, 1985; Baraduc et al., 2013; Desmurget and Turner, 2010; Dudman and Krakauer, 2016; Manohar et al., 2015; Mazzoni et al., 2007; Rueda-Orozco and Robbe, 2015). The SNr microcircuit properties in our model were constrained to match our prior measurements (Brown et al., 2014). Taken together, this suggests that, at least in the context of the skilled forelimb movements, this model has validity. Nonetheless, the model still requires a number of assumptions. Specifically, while we know a fair bit about neural activity in striatum during a skilled forelimb movement (Panigrahi et al., 2015; Yttri and Dudman, 2016), we still do not know the precise relationship between movement correlates in a neuron’s activity and its pattern of downstream connectivity.

The arguments above suggest that if we could compare, within subjects, the effect of systemic reduction of nigral output versus the local suppression of intranigral inhibition experimentally, then we could distinguish between the competing functional models. As described in the introduction, the spatial intermingling of the somata of tonically spiking neurons and the axon collaterals that mediate intranigral inhibition made it essential to use a perturbation strategy that could suppress synaptic transmission without directly interfering with somatic spiking necessary to propagate nigral output to extranigral targets in the thalamus, superior colliculus and brainstem (Dudman and Gerfen, 2015). Recent work suggests that the Gi-coupled DREADD receptor hM4Di (Armbruster et al., 2007) can potently suppress synaptic transmission in some cell-types (Stachniak et al., 2014). Thus, we examined whether synaptic transmission could be suppressed in nigral projection neurons by co-expressing the optogenetic activator Channelrhodopsin-2 (ChR2) and the hM4Di receptor in GABAergic neurons of the SNr (see Methods).

In acute brain slices from the midbrain, we patched GABAergic neurons in the SNr as described previously (Brown et al., 2014) near the site of ChR2::hM4Di co-expression (Fig. 2A). In individual nigral neurons we found that brief pulses of blue light evoked robust inhibitory post-synaptic currents (IPSCs; Fig. 2B-D) as previously reported (Brown et al., 2014). We next found that bath application of CNO, a ligand of the hM4Di receptor, profoundly suppressed evoked synaptic transmission within a few minutes in all cells tested (Fig. 2B-D). To our surprise, however, and in contrast to many other cell types (Armbruster et al., 2007; Stachniak et al., 2014), there was no measurable effect on somatic excitability (Fig. 2E-H). We detected no change in tonic firing rate, nor the input resistance, slope of the FI curve during injected current, or rheobase measured with injected current steps (Fig. 2E-H). Burst spiking evoked by photostimulation of ChR2+ SNr neurons was also unaffected (Fig. 2I-J). Moreover, extracellular recordings of putative SNr GABAergic neurons *in vivo* likewise showed no change in tonic or light-evoked somatic spiking (Fig. 2K-L). Taken together these data indicate that local infusion of CNO into the nigral neurons expressing the hM4Di receptor can selectively suppress synaptic transmission without producing detectable changes in somatic excitability.

**Figure 2.**
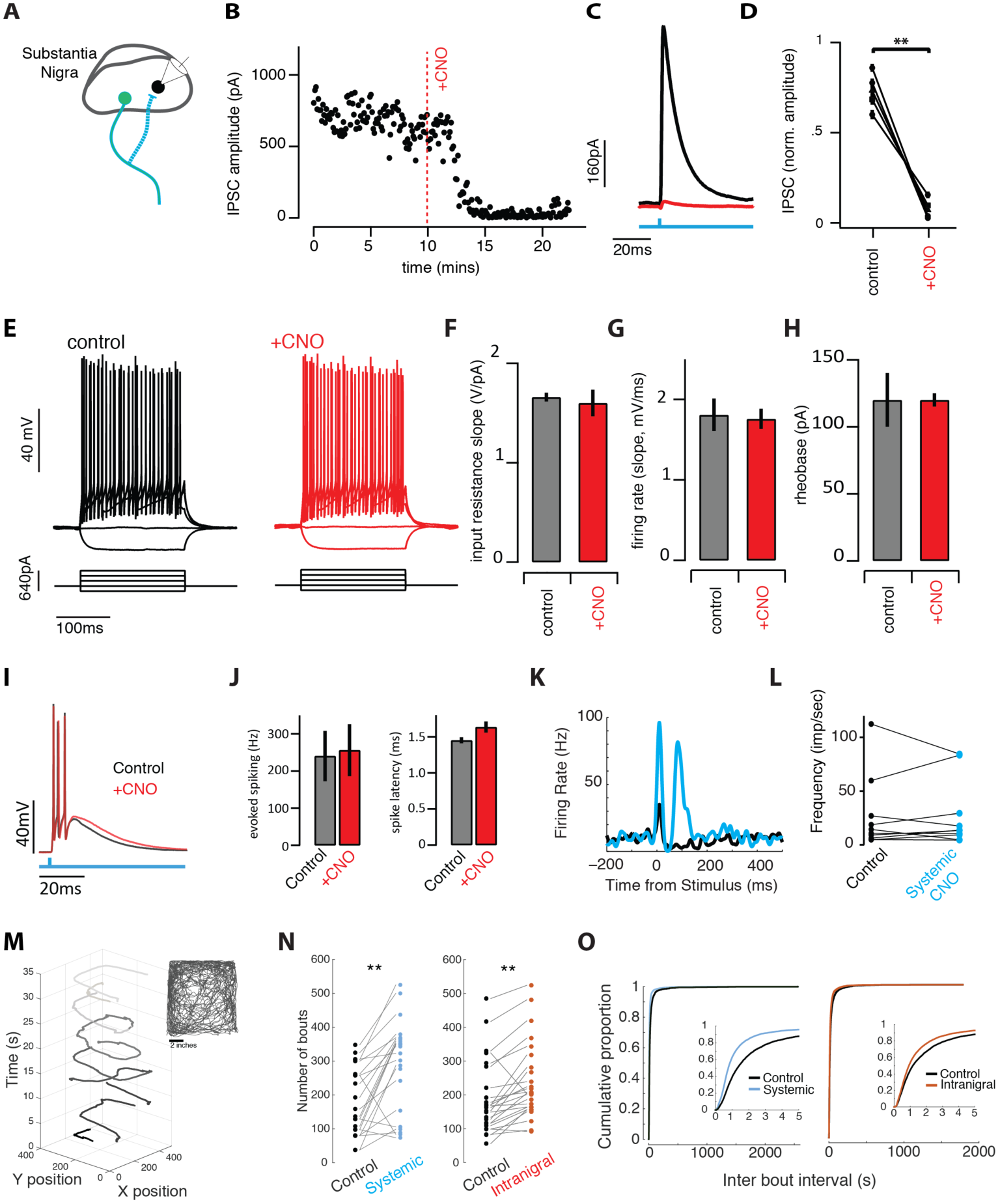
**Suppression of synaptic transmission without changes in excitability allows comparison of intranigral and systemic perturbation of SNr output** **A.** Schematic of experiment. ChR2 and hM4di were co-expressed in GABAergic neurons of the SNr. Intracellular recordings were made from neighboring cells and inhibitory post-synaptic currents (IPSCs) were isolated in voltage clamp. **B.** Light-evoked synaptic transmission was monitored before and after addition of the ligand for the hM4Di receptor, ‘+CNO’, was added to the bath. **C.** Average IPSCs recorded during a baseline period (black) and following infusion of CNO to the bath (red). **D.** Population summary of the suppression by CNO. **E.** Cellular excitability was estimated by holding cells at ∼-70mV and injecting decreasing and increasing current steps. **F-H.** Summary data for a number of parameters extracted from the input-output curves (E) in control (black/gray) and +CNO (red) conditions. **I.** Light-evoked spiking (500ms pulse) evoked burst spiking both (black) before and after (red) perfusion of SNr brain slices with CNO. **J.** Population summary of light-evoked spike frequency and onset latency before and after CNO. **K.** Light evoked spike PSTH of example SNr unit in vivo before (black) and after (blue) systemic CNO. **L.** Population summary of baseline firing rate of single units before and after systemic CNO injection. **M.** The animal’s movement trajectory (seen in inset) can separated into distinct bouts of movement. **N-O.** Systemic and intranigral CNO infusions increased the number of spontaneous bouts of movement and changed the distribution of time between bout initiation, reducing the mean inter bout interval.

We next sought to confirm that systemic and intranigral suppression of GABAergic transmission were sufficient to produce changes in voluntary movement. Adeno-associated viruses (AAV) containing a cre-dependent hM4Di::eGFP transgene were injected into the SN of mice expressing cre-recombinase under the GAD2 promoter (see Methods; Fig. 2 suppl. 1). Cannulae were chronically implanted bilaterally above the SNr for the local delivery of CNO (or vehicle controls). Mice were placed in an open field and locomotor behavior was examined by tracking the centroid position of the mouse. Bouts of locomotion were isolated as described previously (Panigrahi et al., 2015) and analyzed in detail (see Methods).

The tonic output of the SNr has long been proposed to act as an inhibitory break on movement (Chevalier and Deniau, 1990; Chevalier et al., 1985; Hikosaka and Wurtz, 1985). Thus, inactivation of SNr output is predicted to increase the prevalence and vigor of voluntary movement via disinhibition. Consistent with this prediction, systemic suppression of nigral output dramatically increased the number of movement bouts initiated (Fig. 2N) and reduced the inter-bout interval (Fig. 2O). Local, intranigral infusion of CNO produced similar alterations to locomotion (Fig. 2N-O) indicating that suppression of intranigral inhibition can effectively alter SNr activity. The velocity of individual bouts of locomotion and total distance travelled were also increased following systemic CNO; however effects were subtly different following intranigral infusion (Fig. 2 suppl. 2). In both systemic and intranigral sessions, animals initiated more, faster bouts of movement at a higher rate, leading to an increase in total distance traveled following CNO infusion compared with vehicle infusion. We note that alterations in behavior were not associated with any gross coordination changes. The stride and positioning of the paws during bouts of locomotion in the presence of systemic suppression of nigral output were normal (Fig. 2 supp 2). Thus, the increase in the amount and vigor of movement are consistent with a modified gain of motor output (Dudman and Krakauer, 2016) rather than (mal)adaptive changes to disrupted motor control. It is important to note that on the one hand systemic and intranigral infusions of CNO should produce at least somewhat overlapping effects. Systemic CNO infusion will also block intranigral transmission and thus produce a gain effect. This is important because the penetrance of hM4Di expression is not 100%. A gain effect can thus have behavioral consequences mediated via hM4Di- SNr projection neurons that are not blocked during systemic administration. Thus, it is important that there are dissociations between systemic and intranigral administration of CNO; these differences indicate a differential effect of the two mechanisms of suppression.

In summary, we found that systemic and intranigral suppression of SNr synaptic transmission produced behavioral effects that were consistent in sign. Notably, the time between bouts of locomotion (movement initiation) and total amount of locomotion changed in the same direction for both manipulations contrast with predictions of either the tonic inhibition or lateral inhibition models ((Table 1). However, it is more difficult to make quantitative predictions about the consequences of altering basal ganglia output during voluntary locomotion. Although, disruption of basal ganglia signaling similarly alters both skilled forelimb movements and locomotion (Panigrahi et al., 2015). Specifically, it is less well understood how specific patterns of direct and indirect pathway activity are related to the control of locomotion speed (but see (Rueda-Orozco and Robbe, 2015)). Thus, we next turned to the performance of a skilled forelimb movement where our computational model made specific, quantitative predictions.

We trained mice to perform a skilled forelimb movement task (Fig. 3) described previously (Panigrahi et al., 2015). Briefly, head-fixed mice were trained to displace a joystick past an unsignaled threshold in exchange for a small drop of water. Threshold crossings were not signaled to the mouse, but were followed after 1 second delay with reward delivery. The task was self-paced, but a minimum intertrial interval of 3 seconds was imposed between reward delivery and subsequent threshold crossing (Fig. 3A-B). Mice were injected with AAV-DIO-hM4Di into the SNr and implanted with bilateral cannulae prior to training. After a few weeks of training mice learned to adjust movement amplitudes to the block structure of a threshold staircase (Fig. 3B). We then began testing with a randomly ordered, internal matched control design (see Methods) alternating between systemic injection or intranigral infusion of CNO or vehicle to assess model predictions ((Table 1).

**Figure 3.**
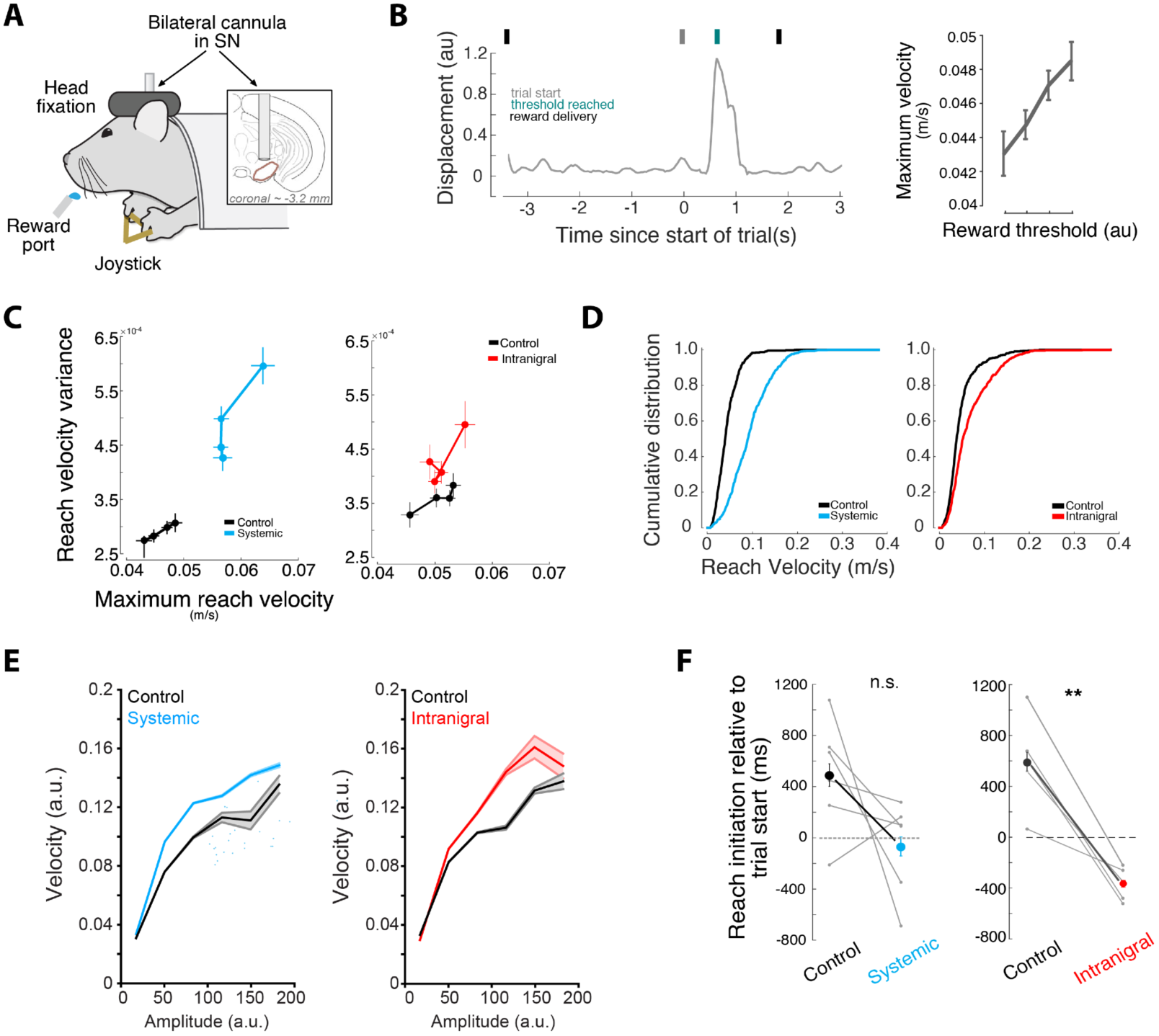
**Behavioral effects of intranigral and systemic perturbation of SNr output in mice trained to perform a skilled forelimb movement task** **A.** Schematic of experiment. Mice were implanted with bilateral cannula targeting the SN. Head-fixed mice were trained to move a joystick to varying thresholds for delayed rewards. **B.** Euclidean displacement of the joystick as a function of time for an example trial. Trial events are indicated by labelled tick marks above. (Right) Population data showing that mice adjust the average movement velocity to the imposed amplitude threshold. **C.** Variance vs. mean peak velocity for trial-completing forelimb movements (‘reach’) in sessions with systemic CNO administration (‘Systemic’, cyan throughout) or sessions with direct infusion of CNO through the cannula (‘Intranigral’, red throughout). **D.** Cumulative distribution of reach velocities for exemplar systemic and intranigral sessions. **E.** Peak velocity plotted as a function of reach amplitude for a corresponding range of reaches across all session conditions (Left, systemic CNO administration; Right, intranigral CNO administration - analogous to Figure 1G) **F.** Time of trial initiation relative to the elapsed intertrial interval (3 seconds) for sessions with systemic (left, cyan) or intranigral (right, red) administration of CNO.

First, we analyzed the average trajectory (Fig. 3 suppl. 1), amplitude (Fig. 3 suppl. 1), and variance of the max velocity for systemic and intranigral sessions (Fig. 3). As expected, systemic suppression of nigral output produced a large increase in the average amplitude of forelimb movements across all blocks (Fig. 3C-D). In contrast, and consistent with our model predictions for a feedback gain control ((Table 1; Fig. 1), we found that intranigral suppression of inhibition increased the variance of the max velocity of movement, but did not alter the average amplitude (Fig. 3C-D). As predicted (Fig. 1G), a change in movement variance associated with movements of a given amplitude produces a change in the relationship between peak velocity and amplitude (Fig. 3E).

We next looked at the timing of trial initiation across conditions. After a few weeks of training, mice were able to pace their initiation of trials accurately to occur just before the ITI had elapsed (Fig. 3F). Reliably initiating trials in the absence of any external, overt cue indicates that mice accurately time the latency from the previous reward. Systemic suppression of SNr output was associated with an observed decrease in the latency between previous reward and the initiation of the subsequent trial, however, this effect was not consistent across all mice or sessions (Fig. 3F). In contrast, intranigral suppression of inhibition was associated with a consistent decrease in the average time between reward and trial initiation that was consistent across all sessions and mice (Fig. 3F). The direction of the change in the timing of trial initiation has the same sign in the two conditions tested (although it is unreliable in the systemic condition) consistent with increased gain during intranigral inhibition and a bias towards reduced activity during systemic suppression.

A large body of literature has implicated the basal ganglia in the timing of actions in the supra-second range (Merchant et al., 2013). There are several proposed mechanisms for timing computations in the basal ganglia. In general, perturbation experiments suggest that activation of D1 receptor-expressing neurons in striatum (and thus putative reduction in firing in SNr) produce decreases in the time between actions (‘the clock running faster’) consistent with our observation that an increase in gain, putatively enhanced suppression of SNr activity, decreases the timed interval between trials (Buhusi and Meck, 2005). However, the reliability of the intranigral manipulation relative to the systemic manipulation suggests that perhaps the contrast between direct and indirect pathway activity might be critical for timed behaviors (Yin, 2014). Specifically, enhanced gain should enhance contrast between direct (suppression) and indirect (activation) in SNr projection neurons. Systemic suppression of output would counteract this contrast by globally reducing output.

## Summary

Here we have shown that a combination of diverse tonic firing rates, a spatially diffuse collateral circuit, and nonlinear subthreshold response to asynchronous inhibition can be sufficient to implement divisive feedback inhibition in a model of the SNr microcircuit. This model incorporates the salient features of the circuit and cellular biophysical properties of the SNr we reported previously (Brown et al., 2014). This model exhibits a form of gain control that can be revealed by selectively manipulating synaptic release within the SNr. By integrating this circuit into a previously described circuit model for the control of forelimb movements (Yttri and Dudman, 2016) we predicted that intranigral suppression of synaptic transmission would produce specific, but subtle changes to behavior ((Table 1) - notably an increase in the variance of movement velocity. More importantly, other proposed functions of intranigral inhibition - lateral inhibition and tonic suppression - made distinct behavioral predictions ((Table 1). Exogenous activation of Gi-coupled signaling pathways can suppress synaptic transmission (Stachniak et al., 2014). Here we show that in SNr projection neurons the suppression of synaptic transmission is, in fact, the primary consequence of activation of Gi-coupled signaling via the hM4Di DREADD receptor and no associated change in somatic excitability was detected *in vitro* or *in vivo*. By comparing local infusion and systemic administration of the hM4Di receptor ligand, CNO, we could thus compare the role of intranigral inhibition with suppression of SNr output to extra-basal ganglia targets.

We performed a comparison on systemic and intranigral perturbations in both freely moving mice in an open arena and in mice trained to perform a skilled forelimb movement for reward (two tasks that require intact basal ganglia signaling (Panigrahi et al., 2015)). We summarize the observed results of our perturbation experiments in (Table 1. The predictions from a feedback gain control functional model and the observed behavioral results match very accurately, whereas other functional models are less consistent with our data. Taken together our data demonstrate that the unique biophysical and circuit properties of the nigral microcircuit are sufficient to implement divisive feedback inhibition which functions like a feedback gain control on the basal ganglia output. Moreover, our perturbation experiments indicate that gain control appears to be the dominant mode in which the SNr operates during voluntary locomotion and purposive, skilled forelimb movements.

Here we show that exogenous modulation of Gi-coupled signaling in SNr projection neurons can increase the variability of movement kinematics and bias the timing of trial initiation. From a broad perspective a change in the gain of the basal ganglia output can be viewed as producing relatively modest changes in behavior. Mice still perform a purposive behavior accurately enough to collect reward (Fig. 3 suppl. 1D) and the basic properties of exploratory behavior (locomotion in an open field environment) appear intact (Fig. 2; Fig. 2 suppl. 2). However, from another perspective the observed changes in behavior are particularly interesting *because* they don’t produce profound disruptions to purposive, voluntary behavior. In dynamic environments, it is important to engage in exploratory changes in behavior that may, in short run, make behavior less adaptive, but exploration is critical in the long run to ensure that changes in the contingencies of the environment can be detected and exploited. In many tasks the timing of behavior (Gallistel and Gibbon, 2000) or movement kinematics (Dudman and Krakauer, 2016; Turner and Desmurget, 2010) are critical to adaptive behavior. Exploration of timing or kinematics should be restricted to changes that don’t eliminate successful task performance. In this sense, we propose that endogenous modulation of intranigral synapses via Gi-coupled signaling may provide a circuit mechanism for controlling the exploration of timing and kinematic dimensions of purposive behavior. In future work, we hope to identify metabotropic signaling in the SNr that could selectively regulate intranigral synaptic transmission and examine its role in adaptive behavior.

## Acknowledgements

Tev Stachniak and Scott Sternson provided critical insight at the start of the project. In addition, we are indebted to the extensive feedback from lab members and our colleagues following presentation of this work at internal seminars at Janelia Research Campus. JB was a graduate scholar in the Cambridge-Janelia Farm Graduate Program when some of this work was performed. JTD is a JRC Group Leader of the Howard Hughes Medical Institute. This work was supported by funding from the Howard Hughes Medical Institute. The authors declare no competing financial interests.

## Materials and Methods

### Animals

Adult transgenic mice expressing cre-recombinase under the control of the glutamic acid decarboxylase 2 gene (GAD2-cre; Stock #010802; Jackson Labs) were used in all experiments. Mice were housed in a temperature and humidity controlled room maintained on a reversed 12 hr light/dark cycle. For in vivo physiology and behavioral experiments, mice were housed individually. Following 1 week of recovery from surgery, 8 mice were put onto water restriction where the water consumption was limited to at least 1ml per day. Mice underwent daily health checks and water restriction was eased if mice fell below 70% of their body weight at the beginning of deprivation. For *in vitro* experiments, mice were group housed (1-5 mice per cage). Mice had ad libitum access to water. All animals were handled in accordance with guidelines approved by the Institutional Animal Care and Use Committee of Janelia Research Campus.

### Viral expression

Two adeno-associated viruses (AAV, serotype 2/1) were used to achieve conditional expression of ChR2 (AAV2/1-SYN-FLEX-ChR2-GFP) and the hM4d receptor (AAV2/1.CAG.FLEX.hm4D:2a:EGFP.reverse). Viruses were produced at the Molecular Biology Shared resource of Janelia Research Campus. Viruses were injected into the substantia nigra (SN) of GAD2-cre mice. Animals were deeply anaesthetised under isoflurane (∼0.5% in O2) and a small craniotomy was made over the SN (from bregma: -3 mm anterior-posterior, 1 mm medial-lateral, -4.2 and -4.4mm from the dura dorso-ventral). A glass pipette was used to pressure inject small volumes of virus at a rate of 23 nL/sec (NanoJect) achieving a total injected volume of ∼100nL of virus per injection depth. Animals were given postoperative administration of butorphenol (0.05 mg/kg) and ketoprofen (5 mg/kg) for at least 2 days post-surgery.

### In vitro electrophysiology

Coronal midbrain slices (300um thick) were cut from adult mice for SNr recordings as descried previously in Brown et al 2014. For optogenetic recordings, areas of high ChR2-YFP expression within SN was used to target recording area. Individual neurons were patched under DIC optics and cell attached spiking was used to classify neurons as putative GABAergic neurons (high firing rates and short action potential waveforms (ref)). Whole cell voltage clamp configuration was used to isolate and characterize both synaptically driven and light driven currents. To isolate inhibitory postsynaptic currents (IPSCs) cells were held at ∼+20mV in the presence of excitatory synaptic blockers (AP5 and NBQX). Analysis of postsynaptic currents was performed using custom written analysis code in Igor Pro (Wavemetrics). Peak current amplitude was measured as the peak synaptic current relative to the baseline holding current preceding each stimulus. Cellular excitability was measured using whole cell current clamp configuration. A series of hyperpolarizing and depolarizing current steps were injected to the cell while holding the cells at ∼70mV. The slope of the linear fit between peak change in membrane potential and injected current was used to measure the input resistance of the cell. Spikes were detected at the threshold of maximum acceleration and the intrinsic excitability of a neuron was measured by calculating the slope of the firing frequency-current (f-I) relationship as increasing positive current steps are injected. The rheobase was defined as the current at which the first spike was evoked.

Wide field optical stimulation was used to drive ChR2 positive cells using a blue LED (473nm, ThorLabs) transmitted through the fluorescence light path of the microscope. For CNO (Anzo Life Sciences) experiments, CNO (10uM) was dilute and bath applied in aCSF to the recording solution.

### In vivo electrophysiology

Recordings were performed using a 64-channel silicon probe array (NeuroNexus Technologies). Electrode arrays were stereotaxically implanted under anesthesia (isoflurane, 1.5-2.5% in O2) targeted to the SN (from bregma: -3.0-4.5 mm anterior-posterior, 0.5-2.9 mm medial-lateral, >3.5mm from the dura dorso-ventral). A 200um core multimodal fiber (Thorlabs) was affixed near the central recording shank of the silicon probe array. The entire array was slowly lowered into the midbrain. Single units were obtained from awake, alter mice. Single cell isolation was performed offline using Offline Sorter (Plexon Technologies) and standard techniques. Analysis of stimulus-evoked responses were calculated and presented using Matlab. For in vivo CNO experiments, baseline evoked activity was recorded following 470nm stimulation over SN, and then subsequently following 20 and 30 mins post CNO IP injection.

### Cannulae

For in vivo intranigral delivery of CNO, bilateral cannulas (length: 3.5 mm; injection tube: -4mm) were implanted above the site of injection of virus medial/lateral +/−1.2 mm; anterior/posterior: -3.2 mm. Dental cement cement was used to anchor the guide cannula to the skull. Dummy cannulae (Plastic One) were inserted to keep the cannula from getting clogged.

### CNO and vehicle infusions

On testing days, mice received either a systemic (IP) or intranigral (cannula directed microinjection) injection of either saline or CNO while under light anesthesia (isoflurane). For all experimental infusions, CNO was diluted in buffered saline. For systemic injections, mice received 0.7 ml of either saline or CNO (3 mg/kg). For intranigral injections, mice received 500 nl total (250 nl bilaterally) of saline or CNO (3 um) with a flow rate of 50 nl/min. Thirty minutes post injection, animals performed the head-fixed operant task for 30 minutes and were then placed in the open field behavioral task for an additional 30 minutes.

### Effort-based operant task

Animals were trained on a head-fixed reaching task as previously described (Osborne and Dudman, 2014; Panigrahi et al., 2015). Mice underwent 12 sessions of initial training prior to either systemic or intranigral injections to ensure animals were near behavioral steady state. Mice were places in a darkened chamber with both paws positioned on a small metal handle attached to a joystick with two degrees of freedom (as seen in Figure 3). Movement was coupled to a variable resistor, whose voltage drop was recorded was recorded at 10kHz, filtered, smoothed and downsampled to 250 Hz resolution.

A session consisted of 120 trials within 7 blocks at 4 difficulty levels (Figure 3B). At the start of each trial, the joystick position was reset to the coordinates 0,0 digitally. Animals were trained to maneuver the joystick to certain displacement thresholds. Displacement thresholds were determined by resistance changes of a potentiometer connected to the joystick, including 10, 15, 25, and 30 units (256 samples distributed over +/− 2.5 V giving approximately 10 mV per unit) that were linearly proportional to the displacement threshold length. After a successful threshold crossing, a sweetened water reward (∼0.05-0.1 mL per trial) was delivered after a short delay (1 second).

Reaches were extracted from calculated velocity traces by using a threshold and several other temporal criteria. To extract parameters of the outward component of the reach, we followed a technique described in detail previously (Gallivan and Chapman, 2014). Briefly, the outward component of the reach was determined by finding the end point of maximum displacement from the starting point of the reach (the most eccentric point on the convex hull that captured the trajectory). The outward component was extracted and the major axis of movement determined by the end point vector. This vector was rotated to π/2 using a matrix transform. This allowed us to define a major axis (“ON axis”) and an orthogonal axis (“OFF axis”) for each reach trajectory. A number of characteristics from the reaches were extracted from the x,y position, velocity and acceleration traces.

### Open field behavioral-task

To assess gross motor output of subjects, an open field arena assembled with a non-reflective, black plastic box with dimensions of 7”x12”x13” was used. A high definition camera (Rocketfish) with an aerial view covering the width of the box was used with custom written tracking software to record the movement of the animals.

Prior to experimental days mice were allowed to explore the box for 4 x 30-minute sessions to aid habituation. The x and y position data of the animals were collected at a sampling rate of 60 Hz. These data were used to reconstruct the animals’ position in the open field over time. From here custom written analysis was used to extract several properties of the movement.

To detect movements over the jitter inherent in the x and y position data a ‘centre change’ method was used which was developed previously in the lab (Panigrahi et al 2015). For a change in the x and y coordinates to be classified as a movement, a threshold 2.4 cm (which is the approximate radius of a mouse head) had to be traversed before the movement is detected and a new centre updated. A series of at least 5 centre changes, taking less than 400 ms per centre change, and with an angle variation less than 1, which equates to angles to angle variation from one centre chance to the next of <∼150 degrees, had to occur to be classified as a ‘bout of movement’. From such bouts, kinematic measurements were obtained.

Once each bout within the session had been isolated, properties of these bouts, such as the total distance (converted from pixels to cm (1 cm = 8.3 pixels), total duration, peak velocity and tortuosity (total distance / Euclidian distance between the start and stop of a progression) and inter bout interval were calculated.

### Computational models

#### Simulation of substantia nigra pars reticulata microcircuit

SNr neurons are tonically active with a sharply nonlinear perithreshold conductance (see Fig. 4 supplement 1C from (Brown et al., 2014)). To match these properties we simulated SNr units as conductance-based integrate-and-fire units with a nonlinear bias current (Fig. 1A) governed by the following parameter distributions (where priyakarthick is a normally distributed random variable; and Φ is a cumulant of a normal distribution; both parameterized by mean, standard deviation):

> spike_reset = -65 mV + priyakarthick(0,2)
>
> spike_threshold = -45 mV +priyakarthick(0,2)
>
> time_constant = 0.25 + priyakarthick(0,0.02)
>
> bias_current_lo(mV) = (2.5 + priyakarthick(0,0.33)) * ( -Φ(-65,3) + 1.05 + Φ(-42,1.5))
>
> bias_current_hi(mV) = (1.25 + priyakarthick(0,0.33)) * ( -Φ(-65,3) + 1.05 + Φ(-42,1.5))
>
> hi:lo = 2:1

Connectivity was random governed by a spatial connection density ϕ(0, 0.25) in units of simulated neurons. With a maximal connection probability of 5% and conductance magnitude defined by:

> W(i,j) = -3.3 ./ (N * 0.05)
>
> τ_intranigral_ = 5 ms

A representative weight matrix (normalized magnitude) is shown at right. Typically, simulations were run with N = 480 SNr units, although qualitatively similar results were obtained with simulations ranging over at least an order of magnitude in cell number.

To assess whether the model produced divisive inhibition simulations were run with a randomly varying, balanced, coherent extrinsic input to most units in the simulation (85-100%; also see Fig. 1). Input/ output functions were determined by estimating at each moment the spike density (smoothed with a gaussian kernel; s.d.=10ms) as a function of the normalized, net input for each simulated unit. Input/ output functions were binned (11 evenly spaced bins) and averaged for the entire population and the contrast between simulations plotted with the standard error of the mean. Penetrance was defined as the fraction of units in which weights were reduced to 2% of their initial value.

#### Simulation of ‘full’ basal ganglia circuit

The basal ganglia circuit simulation involved the following steps.

1. Generate ‘motor commands’

> Natural forelimb movements in mice manipulating a joystick range from having unimodal velocity profiles to having more complex, multimodal velocity profiles (Osborne and Dudman, 2014; Panigrahi et al., 2015; Yttri and Dudman, 2016) and are likely to be composed of varying ratios of direct and indirect pathway projection neurons (Dudman and Krakauer, 2016; Yttri and Dudman, 2016). To capture this variance we ran simulations of 200 movement profiles produced from the following generative function:
>
> The net motor command was the sum of 2 repetitions of the function (each repetition corresponding to input to one of the two pathways). For simplicity we will refer to the velocity profile as a ‘reach’.
>
> 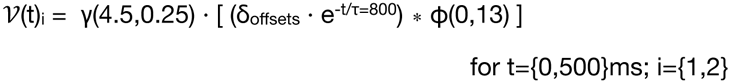
>
> where, ν is velocity; γ is a random variable drawn from a Gamma pdf in arbitrary units of velocity; ϕ is a normal distribution in units of ms; τ is time constant (in ms) reflecting the fall off in velocity (complex movements tend to have early velocity peaks).
>
> The ‘motor command’ is then:
>
> ν = ( ν_indirect_ + ν_direct_) / 2
2. Create a population response in direct and indirect pathway projection neurons of striatum

> Each motor command is used to generate a layer of spike rates corresponding to direct and indirect pathway projection neurons. Poisson spikes are generated according to continuous spike probability functions that are either monotonically tuned to velocity (Yttri and Dudman, 2016), or untuned:
>
> 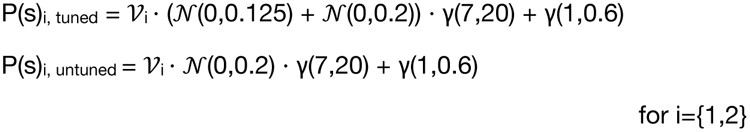
3. Sum the motor command with simulated SNr output

> Step 2 generated spike trains in the striatal layer of the network (see Fig. 1E for an example simulation). Typically, simulations used N_direct_=N_indirect_=5∙N_SNr_. For reported simulation data N_direct_=2400, although simulations were run over an order of magnitude range of N with qualitatively similar results. Spike trains were convolved with exponential post-synaptic current waveforms governed by a decay time constant and an offset (τdirect=12ms; t_off_direct_=0; τ_direct_=12ms; t_off_indirect_=15ms) to generate continuous conductance inputs to the SNr layer. Currents were generated assuming reversal potentials of E_direct_=-70; E_indirect_=0; E_intranigral_=-70.
>
> STR->SNr weights were randomly chosen from connection probability function centered on the unit:
>
> P(directi>SNr) = ϕ(0, 0.15∙N_SNr_)
>
> W(directi>SNr) = 300 / (N_direct_ ∙ 0.25)
>
> P(indirecti>SNr) = ϕ(0, 0.67∙N_SNr_)
>
> W(indirecti>SNr) = 50 / (N_direct_ ∙ 0.25)
4. Integrate the net motor command to produce a predicted ‘forelimb movement’

> To produce a predicted movement trajectory the motor command was combined with the tuned SNr output population (typically F_tuned_=0.4) projected on to a ‘decoding’ vector with random weights distributed uniformly over {0,1}.
>
> Y(t) = ∫ ( ν - 0.2[ d(1,N_SNr_∙F_tuned_) ∙ r(N_SNr_∙F_tuned_,t) ])

### Statistics

All statistical tests were performed using the statistics package from Matlab 2014b (Mathworks). Paired comparisons were performed using the student’s t test. Multiple comparisons were performed using ANOVA. Cumulative distributions were compared using a two-sample Kolmogorov-Smirnov test. Significance was defined as p < 0.05 unless otherwise indicated. Averaged data are presented as mean ± standard error of the mean (SEM), unless otherwise specified.

| Subplot | Description of plot | Systemic | Intranigral |
| --- | --- | --- | --- |
| D | Paired comparison of IPSC amplitude before and after CNO | N/A | paired t-test; $p < 0.01$ |
| ALL | All other comparisons were paired t-tests that were non-significant |  |  |

| Subplot | Description of plot | Systemic | Intranigral |
| --- | --- | --- | --- |
| N | Paired comparison of number of spontaneous bouts during locomotion - comparison to a within week control | paired t-test; $p < 0.01$ | paired t-test; $p < 0.01$ |
| O | cumulative distribution of inter-bout intervals - inset shows 0-5 s IBI | two-sample Kolmogorov-Smirnov; $p = 1.8145 \times 10^{-126}$ | two-sample Kolmogorov-Smirnov; $p = 3.3876 \times 10^{-46}$ |

| Subplot | Description of plot | Systemic | Intranigral |
| --- | --- | --- | --- |
| C | blockwise means of velocity vs velocity variance for all rewarded reaches in block | one-way ANOVA; velocity ($p=0.0012$) velocity variance ($p = 0.002$) | one-way ANOVA; velocity ($p = 0.6822$); velocity variance ($p = 0.0309$) |
| D | cumulative distribution of rewarded reach velocities for one animal, all sessions | two-sample Kolmogorov-Smirnov; $p = 3.47 \times 10^{-20}$ | two-sample Kolmogorov-Smirnov; $p = 2.463 \times 10^{-29}$ |
| F | Mean time of trial initiation - each point represents one animal, all reaches | paired t-test; $p = 0.130$ | paired t-test; $p < 0.05$ |

---

**Figure 2 supplement 1.**
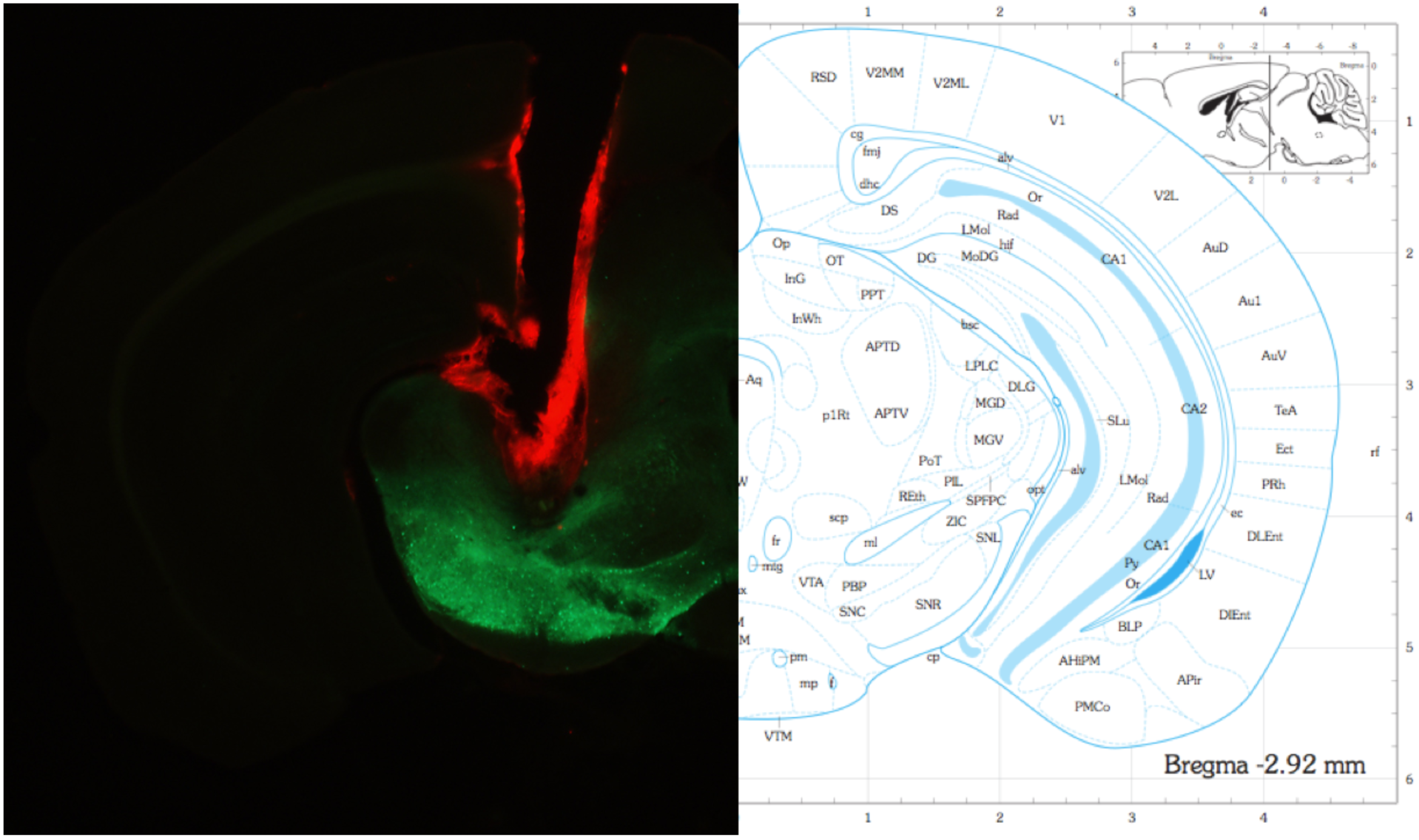
(Left) Example of one hemisphere cannula placement dorsal to the substantia nigra. Cannula was stained with DiI to aid visualization. Note: injection needle extended ∼0.5mm below the cannula for infusions. (Right) Corresponding section from the mouse brain atlas (Paxinos and Franklin, 2004).

**Figure 2 supplement 2.**
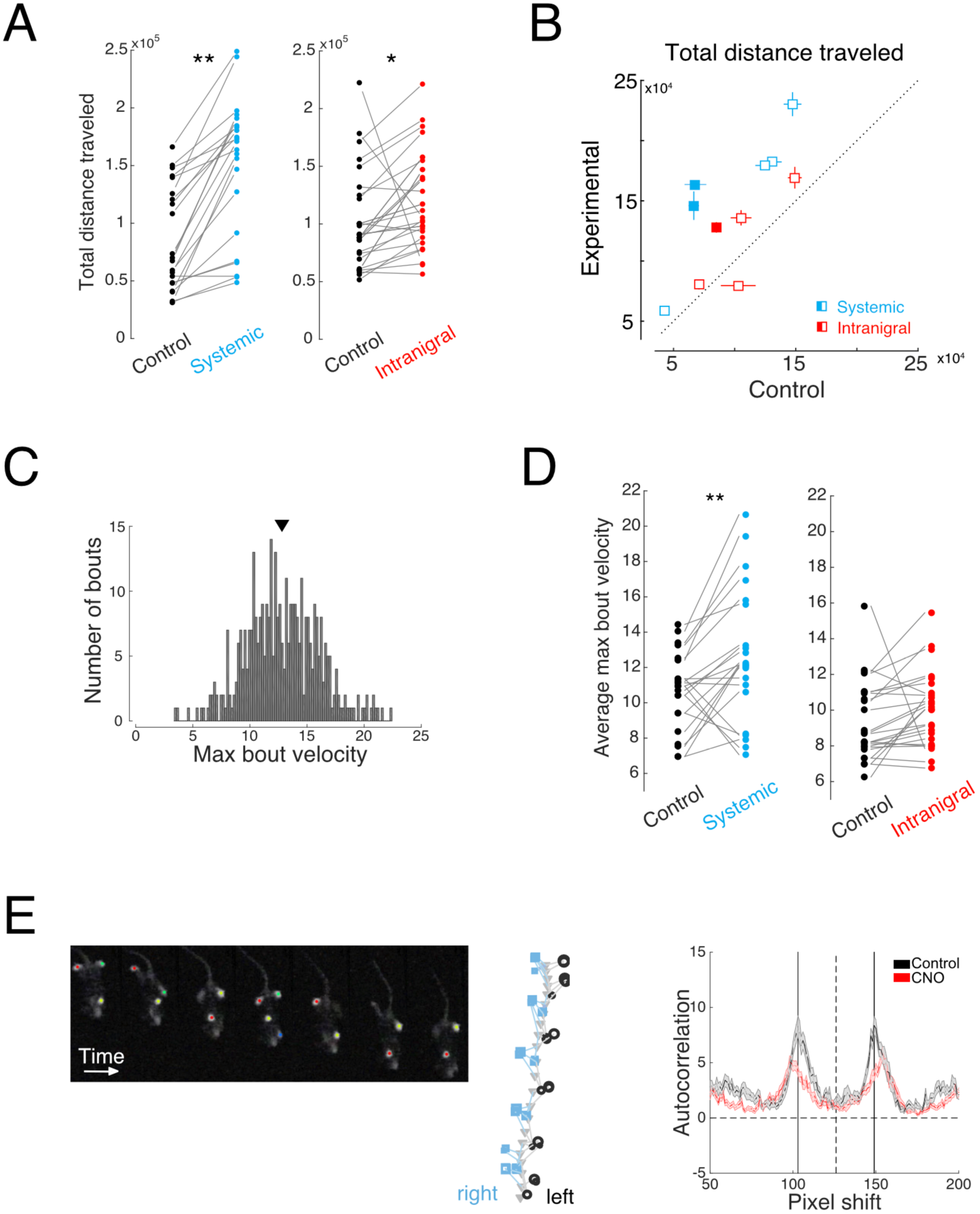
**A.** Both systemic and intranigral CNO infusions increase the total distance traveled in a session (paired t-test, systemic: p < 0.01, intranigral: p = 0.0163). Each point represents the total distance traveled during one 30 minute session with each pair of connected points representing the experimental and control session in one week. **B.**Individual animals show variability, but generally increased distance traveled per session with both systemic and intranigral CNO applications. Each point represents the mean total distance traveled for one animal in control sessions v experimental sessions for at least 2 sessions in each condition. Open squares indicates the animal did not reach statistical significance (paired t-test, p<0.05) **C.** Example distribution of maximum bout velocity from one animal. The black triangle representsthe mean of the distribution. **D.** Systemic (blue), but not intranigral (red), infusion of CNO increases the average maximum bout velocity (paired t-test, systemic: p = 0.0056, intranigral: p = 0.1439). **E.** Example of paw tracking to test that stride and positioning of paws were normal.

**Figure 3 supplement 1.**
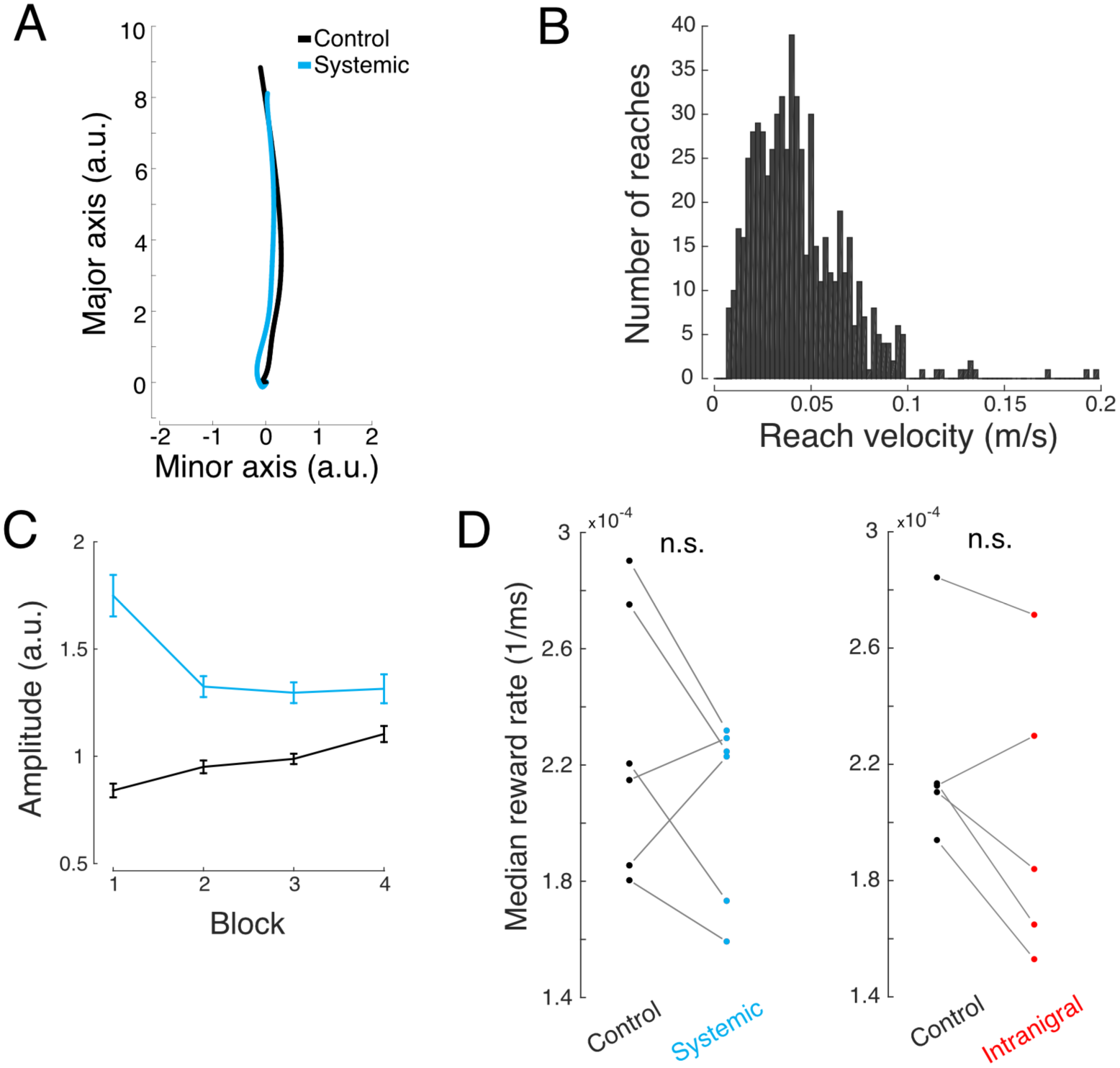
**A.** Example reach trajectories for one animal in control (black) and systemic CNO (cyan) signals aligned to the major axis as described in Gallivan and Chapman (2014). **B.** Distribution of reach velocities for control sessions for the same animal in A. **C.** Mean amplitude per block in control (black) and systemic CNO (cyan) sessions for the same animal as in A. D. Reward rate was not significantly changed in system\ic (cyan) and intranigral (red) conditions in comparison to controls for all animals.

## Footnotes

1 “If the output of MPGs [motor programs] involved in maintaining the body upright against gravity cannot be inhibited for the reaching arm, the voluntary movement of that arm will meet resistance from mechanisms that are trying to keep it in place. The reach may still be accomplished, but only slowly and with great effort since the voluntary activation of agonists in the reach would have to overcome the postural activity of antagonists.”

## Bibliography

Anderson, M.E., and Horak, F.B. (1985). Influence of the globus pallidus on arm movements in monkeys. III. Timing of movement-related information. Journal of neurophysiology 54, 433–448.

Armbruster, B.N., Li, X., Pausch, M.H., Herlitze, S., and Roth, B.L. (2007). Evolving the lock to fit the key to create a family of G protein-coupled receptors potently activated by an inert ligand. Proc Natl Acad Sci U S A 104, 5163–5168.

Bar-Gad, I., Morris, G., and Bergman, H. (2003). Information processing, dimensionality reduction and reinforcement learning in the basal ganglia. Prog Neurobiol 71, 439–473.

Baraduc, P., Thobois, S., Gan, J., Broussolle, E., and Desmurget, M. (2013). A common optimization principle for motor execution in healthy subjects and parkinsonian patients. The Journal of neuroscience: the official journal of the Society for Neuroscience 33, 665–677.

Bays, P.M., and Wolpert, D.M. (2006). Computational principles of sensorimotor control that minimize uncertainty and variability. In The Journal of Physiology, pp. 387–396.

Brown, J., Pan, W.X., and Dudman, J.T. (2014). The inhibitory microcircuit of the substantia nigra provides feedback gain control of the basal ganglia output. Elife 3, e02397.

Buhusi, C.V., and Meck, W.H. (2005). What makes us tick? Functional and neural mechanisms of interval timing. Nat Rev Neurosci 6, 755–765.

Chevalier, G., and Deniau, J.M. (1990). Disinhibition as a basic process in the expression of striatal functions. Trends in Neurosciences 13, 277–280.

Chevalier, G., Vacher, S., Deniau, J.M., and Desban, M. (1985). Disinhibition as a basic process in the expression of striatal functions. I. The striato-nigral influence on tecto-spinal/tecto-diencephalic neurons. Brain Research 334, 215–226.

Cui, G., Jun, S.B., Jin, X., Pham, M.D., Vogel, S.S., Lovinger, D.M., and Costa, R.M. (2013). Concurrent activation of striatal direct and indirect pathways during action initiation. Nature 494, 238–242.

Deniau, J.M., Mailly, P., Maurice, N., and Charpier, S. (2007). The pars reticulata of the substantia nigra: a window to basal ganglia output. Prog Brain Res 160, 151–172.

Desmurget, M., and Turner, R.S. (2010). Motor sequences and the basal ganglia: kinematics, not habits. Journal of Neuroscience 30, 7685–7690.

Dudman, J.T., and Gerfen, C.R. (2015). The Basal Ganglia. In The Rat Nervous System, G. Paxinos, ed. (Amsterdam: Elsevier).

Dudman, J.T., and Krakauer, J.W. (2016). The basal ganglia: from motor commands to the control of vigor. Curr Opin Neurobiol 37, 158–166.

Gallistel, C.R., and Gibbon, J. (2000). Time, rate, and conditioning. Psychological review 107, 289–344.

Gallivan, J.P., and Chapman, C.S. (2014). Three-dimensional reach trajectories as a probe of real-time decision-making between multiple competing targets. Frontiers in neuroscience 8, 215.

Hikosaka, O., and Wurtz, R. (1985). Modification of saccadic eye movements by GABA-related substances. II. Effects of muscimol in monkey substantia nigra pars reticulata. In Journal of neurophysiology, pp. 292–308.

Horak, F.B., and Anderson, M.E. (1984). Influence of globus pallidus on arm movements in monkeys. II. Effects of stimulation. Journal of neurophysiology 52, 305–322.

Humphries, M.D., Stewart, R.D., and Gurney, K.N. (2006). A physiologically plausible model of action selection and oscillatory activity in the basal ganglia. The Journal of neuroscience: the official journal of the Society for Neuroscience 26, 12921–12942.

Isaacson, J.S., and Scanziani, M. (2011). How inhibition shapes cortical activity. Neuron 72, 231–243.

Manohar, S.G., Chong, T.T., Apps, M.A., Batla, A., Stamelou, M., Jarman, P.R., Bhatia, K.P., and Husain, M. (2015). Reward Pays the Cost of Noise Reduction in Motor and Cognitive Control. Curr Biol 25, 1707–1716.

Mazzoni, P., Hristova, A., and Krakauer, J.W. (2007). Why don’t we move faster? Parkinson’s disease, movement vigor, and implicit motivation. The Journal of neuroscience: the official journal of the Society for Neuroscience 27, 7105–7116.

Merchant, H., Harrington, D.L., and Meck, W.H. (2013). Neural basis of the perception and estimation of time. Annual review of neuroscience 36, 313–336.

Mink, J.W. (1996). The Basal Ganglia: Focused selection and inhibition of competing motor programs. Progress in Neurobiology 50, 381–425.

Oorschot, D.E. (1996). Total number of neurons in the neostriatal, pallidal, subthalamic, and substantia nigral nuclei of the rat basal ganglia: a stereological study using the cavalieri and optical disector methods. The Journal of Comparative Neurology 366, 580–599.

Osborne, J.E., and Dudman, J.T. (2014). RIVETS: a mechanical system for in vivo and in vitro electrophysiology and imaging. PloS one 9, e89007.

Pan, W.X., Brown, J., and Dudman, J.T. (2013). Neural signals of extinction in the inhibitory microcircuit of the ventral midbrain. Nat Neurosci 16, 71–78.

Panigrahi, B., Martin, K.A., Li, Y., Graves, A.R., Vollmer, A., Olson, L., Mensh, B.D., Karpova, A.Y., and Dudman, J.T. (2015). Dopamine Is Required for the Neural Representation and Control of Movement Vigor. Cell 162, 1418–1430.

Rueda-Orozco, P.E., and Robbe, D. (2015). The striatum multiplexes contextual and kinematic information to constrain motor habits execution. Nature neuroscience 18, 453–460.

Silver, R.A. (2010). Neuronal arithmetic. Nat Rev Neurosci 11, 474–489.

Stachniak, T.J., Ghosh, A., and Sternson, S.M. (2014). Chemogenetic synaptic silencing of neural circuits localizes a hypothalamus-->midbrain pathway for feeding behavior. Neuron 82, 797–808.

Turner, R.S., and Desmurget, M. (2010). Basal ganglia contributions to motor control: a vigorous tutor. Curr Opin Neurobiol 20, 704–716.

Yin, H.H. (2014). Action, time and the basal ganglia. Philos Trans R Soc Lond B Biol Sci 369, 20120473.

Yttri, E.A., and Dudman, J.T. (2016). Opponent and bidirectional selection of movement parameters in the basal ganglia. Nature 533, 402–406.

